# Basal forebrain parvalbumin neurons mediate arousals from sleep induced by hypercarbia or auditory stimuli

**DOI:** 10.1101/766659

**Authors:** James T. McKenna, Stephen Thankachan, David S Uygun, Charu Shukla, Joshua Cordeira, James M. McNally, Fumi Katsuki, Janneke Zant, Mackenzie C. Gamble, Karl Deisseroth, Robert W. McCarley, Ritchie E. Brown, Robert E. Strecker, Radhika Basheer

## Abstract

Brief arousals from sleep in patients with sleep apnea and other disorders prevent restful sleep, and contribute to cognitive, metabolic and physiologic dysfunction. Little is currently known about which neural systems mediate these brief arousals, hindering the development of treatments. The basal forebrain (BF) receives inputs from many nuclei of the ascending arousal system. These inputs include the brainstem parabrachial neurons which promote arousal in response to elevated blood carbon dioxide levels, as seen in sleep apnea. Optical inhibition of the terminals of parabrachial neurons in the BF impairs cortical arousals to hypercarbia, but which cell types within the BF mediate cortical arousals in response to hypercarbia or other sensory stimuli is unknown. Here using optogenetic techniques in mice, we show that BF parvalbumin (PV) neurons fulfill several criteria for a system mediating brief arousals from sleep. Optical stimulation of BF PV neurons during the light period, when mice normally sleep, caused rapid transitions to wakefulness and increased wake bout durations. Unlike many other ascending arousal systems, arousals induced by stimulation of BF PV neurons were brief, resulting in only a small (13.6%) increase in the total amount of wakefulness. Bilateral optical inhibition of BF PV neurons increased the latency to arousal produced by hypercarbia or auditory stimuli. Thus, BF PV neurons are an important component of the brain circuitry which generates brief arousals from sleep in response to internal and external sensory stimuli.

## INTRODUCTION

Sleep is a reversible state of unconsciousness which promotes optimal function of the brain and body. The reversibility of sleep is important since the reduced awareness of the environment and internal milieu is potentially harmful for the animal. Thus, mechanisms exist to rapidly rouse animals from sleep in the presence of stimuli which indicate danger.

Electroencephalographam (EEG) recordings in humans and animals show that in normal individuals brief cortical arousals from sleep occur throughout the sleep period (Halasz et al., 2004; Prerau et al., 2017). The frequency of these arousals is markedly increased in individuals with sleep disorders such as sleep apnea (Guilleminault et al., 1976) and insomnia (Wei et al., 2017), as well as in aged individuals (Li et al., 2018). Elevated levels of sleep fragmentation interfere with the restorative effects of sleep (McKenna et al., 2007) and has numerous damaging consequences, including impaired synaptic plasticity and memory formation (Tartar et al., 2006), disrupted cardiovascular regulation (Somers et al., 2008) and immune system dysfunction (Luyster et al., 2012; Besedovsky et al., 2019).

Considerable work over the past hundred years has identified multiple interlocking arousal systems, collectively known as the ascending reticular activating system (ARAS; Brown et al., 2012). However, which of these systems is involved in brief arousals from sleep is poorly understood. One important brain region which may be involved is the basal forebrain (BF). The BF is considered the final node of the ventral arm of the ARAS, since it receives input from multiple arousal systems in the brainstem and hypothalamus and contains several neurotransmitter systems with projections to the cortex (Jones, 2003; Zaborszky et al., 2012; Brown et al., 2012; Lin et al., 2015; Yang et al., 2017). Recent studies demonstrated that brainstem parabrachial nucleus glutamatergic neurons relay signals about blood carbon dioxide (CO2) levels and mediate arousals from sleep in response to hypercarbia (excessive CO2) (Kaur et al., 2013, 2017; Kaur and Saper 2019). These neurons project to the BF, and optogenetic inhibition of their terminals in the BF markedly delayed arousal from sleep in response to hypercarbia (Kaur 2017). However, which specific type of BF neurons are involved in brief arousals from sleep in response to hypercarbia and other sensory stimuli is unknown.

The BF is made up of three major neuron populations, broadly defined by their neurotransmitter. The best-known neurotransmitter system within BF is the cholinergic system. However, the BF GABAergic system is the largest, and a major subset of the GABAergic neurons express parvalbumin (PV). Previous studies showed that although stimulation of cholinergic neurons can cause arousals from sleep (Zant et al., 2016; Irmak and de Lecea 2014), the response latency is rather slow and requires local interactions with other neurons in the BF, as shown with local BF perfusion of acetylcholine antagonists (Zant et al., 2016; Yang et al., 2014). Thus, cholinergic neurons are unlikely to be a primary mediator of the brief arousals from sleep such as seen in sleep apnea. Here we focus on BF PV neurons, which have an electrophysiological firing profile which suggests a major role in cortical activation and, possibly, promotion of rapid arousals (Duque et al., 2000; Kim et al., 2015; Xu et al., 2015). *In vitro* recordings of identified BF neurons confirmed that BF PV neurons fired at high frequencies (∼20-60 Hz; McKenna et al., 2013), similar to the rapid firing observed in recordings of *in vivo* optogenetically-identified BF PV neurons (Kim et al., 2015). Furthermore, optogenetic excitation of BF PV neurons produced an increase in largely desynchronized cortical EEG activity indicative of wakefulness (Kim et al., 2015; Xu et al., 2015).

To be effective in mediating arousal responses to elevated blood CO2 (hypercarbia) or other important sensory stimuli, an arousal system should ideally generate a rapid cortical response, in preparation for a more prolonged period of wakefulness if the threat persists. However, to not disrupt sleep more than necessary, these arousals should be brief. Thus, in our first set of experiments, we used optogenetic techniques in mice to precisely measure the latency to arousal from sleep, as well as the overall effect on total amounts of sleep, in response to bilateral optical stimulation of BF PV neurons. Subsequently, we used bilateral optical inhibition to directly test whether BF PV neurons mediate arousals from sleep in response to hypercarbia or auditory stimuli. Optogenetic stimulation of BF PV neurons led to rapid cortical arousals but these arousals were mostly brief, resulting in minor changes in total amount of wakefulness and slightly increased wake bout durations, reminiscent of the disruptions of sleep observed in sleep apnea and other sleep disorders. Furthermore, optogenetic inhibition of BF PV neurons delayed or prevented arousals from sleep produced by hypercarbia or auditory stimuli. Thus, BF PV neurons exhibit several properties consistent with a role in mediating brief arousals from sleep in response to hypercarbia and other sensory stimuli.

## RESULTS

### BF PV optogenetic stimulation significantly decreased the median latency for transitions from NREM sleep to wake

BF PV neurons were bilaterally transduced with a channelrhodopsin2 (ChR2) and enhanced YFP (EYFP) fusion protein expressing adeno-associated virus (AAV) in PV-Cre mice (**Figure 1A**; Sohal et al., 2009; Cardin et al., 2009; Kim et al., 2015). At least one month later, transduced BF PV neurons were optically stimulated bilaterally with 40 Hz trains of 10 ms light pulses (473-nm) for 5 s, followed by 55 s of no stimulation. This stimulation cycle was repeated for six hours during the light period when mice prefer to sleep (**Figure 1B**). The stimulation frequency of 40 Hz is within the natural discharge range of BF PV neurons during brain activated states (Kim et al., 2015; Xu et al., 2015), and was chosen since we previously found that this frequency of stimulation was particularly effective in activating the cortex (Kim et al., 2015). Mock stimulation (laser off) was performed on a different day, counterbalanced with the stimulation days.

**Figure 1.**
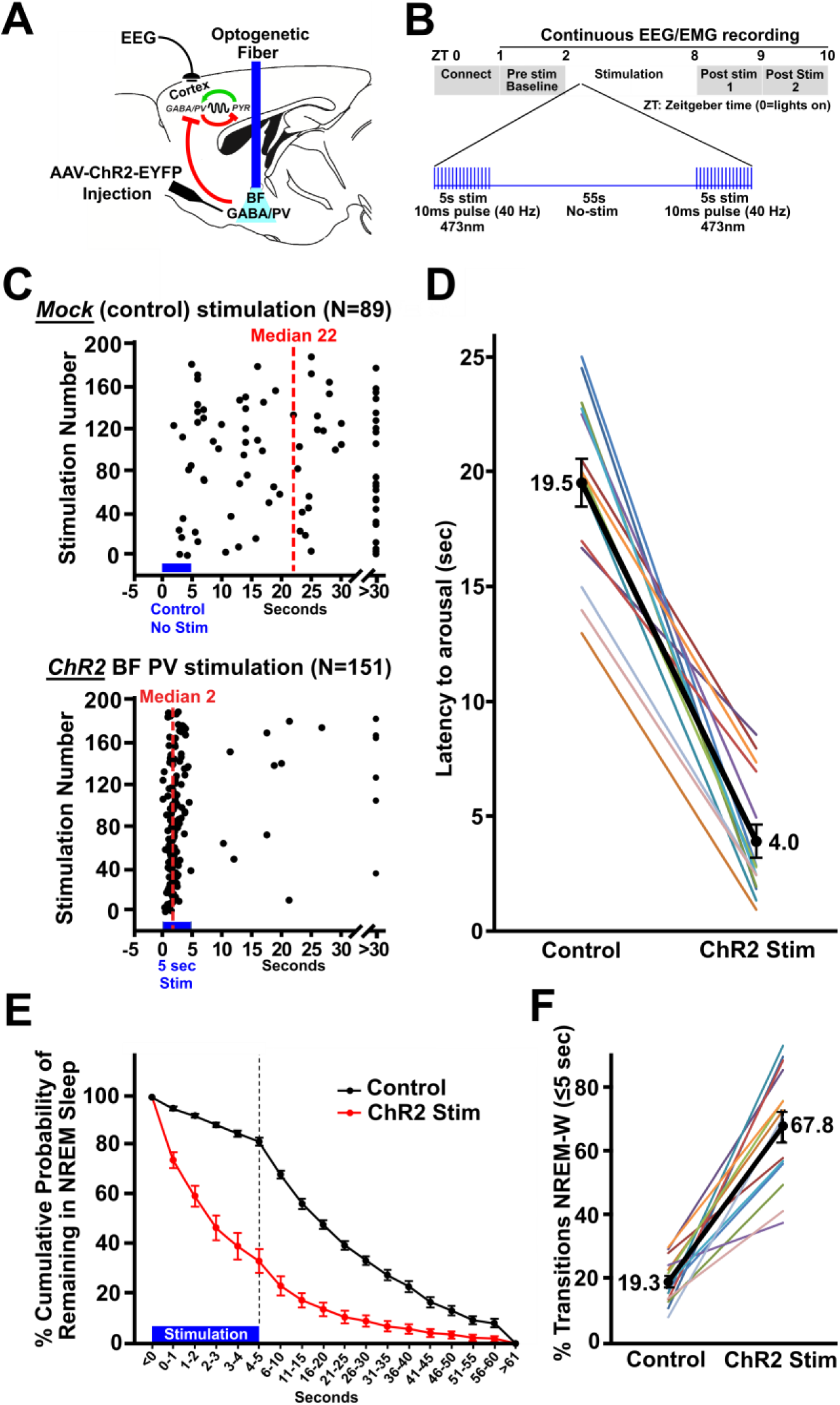
Bilateral optogenetic stimulation of basal forebrain (BF) parvalbumin (PV) neurons strongly reduced the latency to arousal from non-REM sleep. (A) Adeno-associated viral vectors with Cre-dependent expression of channelrhodopsin2-enhanced yellow fluorescent proteins (AAV-DIO-ChR2-EYFP) were bilaterally injected into the BF of PV-cre mice, and optogenetic stimulation applied to elicit arousals. (B) Diagram illustrating the optogenetic stimulation protocol. 40 Hz stimuli with a 10 ms pulse duration was applied simultaneously to both sides of BF for 5 s, followed by 55 s of no stimulation, repeated for 6 hours (ZT2-8). (C) Representative raster plots for one mouse illustrate a profound decrease in the latency for arousal from NREM sleep when stimulation was applied for 6 hours (ZT2-8). (D) Group data for all mice show that optogenetic stimulation produced a significant decrease (∼4 fold) in median NREM-Wake latencies when compared to mock stimulation (control) in the same mice (N=14). (E) The cumulative probability distribution of NREM-to-wake transition latencies shows an increased probability to wake due to ChR2 stimulation compared with Control. The probability of mice remaining in NREM sleep 5 s after the end of stimulation was 33.7%, in contrast to a probability of 82.0% at the same time point in control (mock stimulation). The first 5 s are depicted per second, to demonstrate NREM probability distribution during Chr2 stimulation. 5 s bins are then depicted for the following 55 s. (F) When considering all arousals, the majority (67.8%) of events occurred within 5 s when mice received bilateral stimulation of BF PV neurons, whereas in controls only 19.3% of arousals occurred within 5 s i.e the majority of NREM-W transitions (arousals) occurred within 5 seconds from the start of bilateral BF PV optical stimulation.

Since stimulations were performed at regular intervals, they could occur in any behavioral state. Analysis of those stimulations which occurred during NREM sleep revealed that the latencies to arousal (NREM→wake transition) were significantly decreased when compared to mock stimulation. Raster plot depiction (**Figure 1C**) of one representative case illustrates the striking decrease in the median latency to arousal due to optogenetic stimulation of BF PV neurons. Overall, optogenetic stimulation produced a significant (∼4 fold) decrease in arousal latency (comparison of the means of the median values per animal; Mock Stim Control 19.5±1.0 s vs. ChR2 BF PV Stim 4.0±0.7 s; t=12.123 (df13), P<0.001). We note that this decrease was seen in all mice tested (**Figure 1D**, N=14).

These initial results demonstrated that excitation of BF PV neurons reduced the median latency of arousals from NREM sleep. However, these average data do not indicate whether this is an all-or-nothing effect or reflect a shift in the probability of a NREM-to-wake transition. Probability curve analysis showed a strong drop in the probability that the mice remained in NREM sleep as time progressed per cycle when comparing ChR2 stimulation of BF PV neurons with mock stimulation (**Figure 1E**). The probability of remaining in NREM sleep during ChR2 stimulation trials was significantly decreased compared to mock stimulation (5 s bins analyzed; f=75.16 (df1), P<0.001). By the end of the 5 s stimulation period, the probability of the mouse remaining in NREM sleep dropped to 33.7±4.9 % in mice receiving bilateral stimulation of BF PV neurons, in contrast to 82.0±1.6 % during mock stimulation. The percentage of arousals that occurred within the 5 s of the ChR2 stimulation period was significantly elevated, when compared to mock stimulation (**Figure 1F**; Control 19.3±1.88% vs. ChR2 Stim 67.76±4.83%; t=-9.206 (df13), P<0.001). BF PV stimulation therfore increased the probability of NREM-to-wake transtions, compared to mock stimulation.

### Optogenetic stimulation produced a small increase in the total amount of wakefulness

The amounts of wakefulness, NREM, and REM sleep were evaluated during the six hours of cycling (5s on; 55s off) BF PV optogenetic stimulation (**Figure 2A**). Total amounts of wake were significantly, but only slightly, elevated (Control 36.9±1.2% of total time vs. ChR2 Stim 41.9±1.2%; t= -4.416 (df13), P<0.001; 13.6% increase). NREM sleep amounts were slightly but significantly decreased (Control 54.0±1.2% vs. ChR2 Stim 50.1±1.7%; t= 3.411 (df13), P<0.01; 7.2% decrease), and REM sleep amounts were not significantly altered. As depicted (**Figure 2B**), wake bout durations over the 6 hrs of stimulation were significantly increased due to ChR2 stimulation (Control 49.5±4.5 s vs. ChR2 Stim 59.6±5.9 s; t= -2.326 (df13), P=0.037; +20.5%), and NREM sleep bout durations exhibited a trend-level decrease (Control 68.3±7.1 s vs. ChR2 Stim 58.4±5.2 s; t= 1.958 (df13), P=0.072; -16.2%). Although REM sleep differences did not approach significance, we note that the -17.2% decrease in REM sleep bout durations (Control 71.2±9.6 s vs. ChR2 Stim 58.9±5.2 s) would be predicted to accompany this increase in wake bout durations produced by ChR2 stimulation. To summarize, our stimulation protocol produced rapid arousals from NREM sleep, and the resultant wake bout durations were significantly increased. Overall amounts of wake were not profoundly elevated, though, possibly due to the fact that stimulation was applied regardless of vigilance state, and noting the increased probability that animals were awake when exposed to Chr2 stimulation compared to control (mock) stimulation (**Figure 1E**).

**Figure 2.**
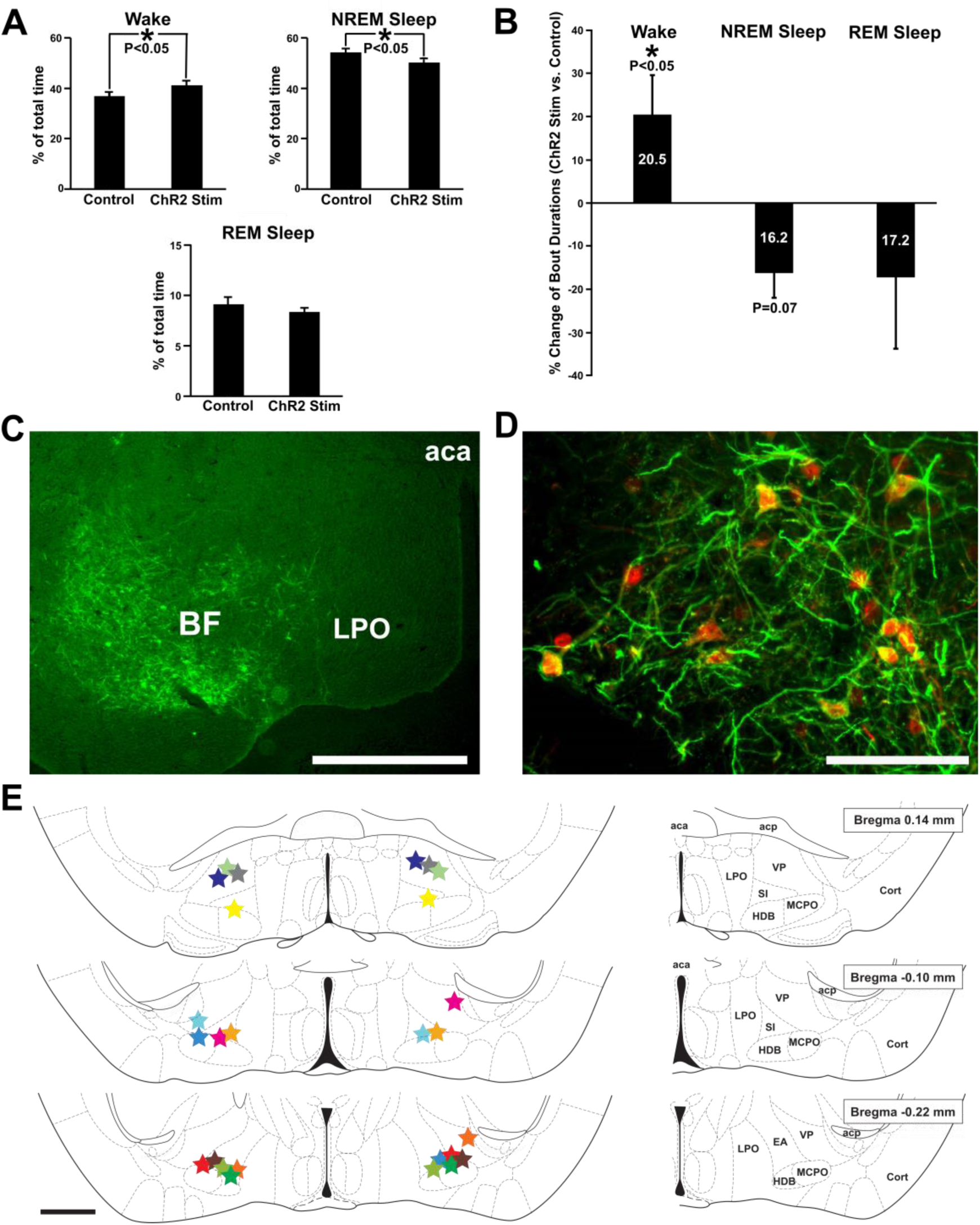
Bilateral optogenetic stimulation of BF PV neurons caused a small but significant increase in the overall amounts of wakefulness and decreased NREM sleep, with successful AAV-ChR2-EYFP transduction and optogenetic probe location verified. **(A)** During the optical stimulation period, total amounts of wake were slightly elevated, whereas NREM sleep amounts decreased, compared to mock (no stimulation) control values. REM sleep was not significantly affected. **(B)** Average wake durations were significantly increased (+20.5%), and NREM sleep showed a trend of decrease (-16.2%), during the stimulation period, compared to control (mock) stimulation. REM sleep diferences were not significant, but indicated a predicted decrease (-17.2%) accompanying the accentuation of wake durations. **(C)** Transduction was evident throughout the majority of BF in all cases targeted for optogenetic stimulation. Arrowhead denotes approximate location of the ventral tip of the fiber optic cannula for one representative case, where blue light stimulation was applied. Scale bar = 1 mm. **(D)** Anti-PV immunohistochemistry (red) shows AAV-ChR2-EYFP injections (amplified with anti-GFP primary antibody; green) selectively transduced PV somata in BF (yellow/orange). As previously reported by our group (Kim et al., 2015), >80% of transduced neurons stained positively for PV. Scale bar = 100 µm. **(E)** Localization of bilateral optogenetic fiber stimulation sites in BF (N=13). Color coding indicates each individual case. Scale bar = 1mm. Abbreviations: aca, anterior aspect of the anterior commissure; acp, posterior aspect of the anterior commissure; BF, basal forebrain; Cort, cortex; EA, extended amygdala; HDB, horizontal limb of the diagonal band; LPO, lateral preoptic nucleus; MCPO, magnocellular preoptic nucleus; SI, substantia innominata; VP, ventral pallidum. Atlas templates schematics adapted from Paxinos and Franklin, 2013.

Histological and immunohistochemical analysis validated the effective and selective ChR2-EYFP transduction of BF PV neurons, as in our previous studies targeting these neurons (Kim et al., 2015; Thankachan et al., 2019; Hwang et al., 2019). AAV-ChR2-EYFP injections transduced neurons and fibers throughout large areas of BF (depicted for one representative case in **Figure 2C**). Transduced neurons were PV-positive, as determined by immunohistochemistry (**Figure 2D**), and similar to that previously reported (Kim et al., 2015). **Figure 2E** depicts the fiber optic cannula placement within BF. The ventral tips of the fiber optic cannula, which are presumably the dorsal-most tips of the area of optogenetic stimulation, were located within BF in 13 cases.

### Optogenetic inhibition of BF PV neurons inhibits arousals due to hypercarbia

In the next series of experiments, we used optogenetic inhibition with ArchT to test whether BF PV neurons play a role in the arousals from sleep induced by internal or external sensory stimuli. AAV-ArchT-GFP was bilaterally injected into the BF of PV-Cre mice, and optical fibers implanted for ArchT activation (**Figure 3A**). Histological and immunohistochemical analysis validated ArchT-GFP transduction of BF PV neurons, similar to our previous studies targeting these neurons (Kim et al., 2015). AAV-ArchT-GFP injections transduced neurons and fibers throughout large areas of BF (depicted for one representative case in **Figure 3C**). The ventral tips of the fiber optic cannula, indicating the dorsal-most tips of the area of optogenetic stimulation, were located within BF in 6 cases (**Figure 3D**).

**Figure 3.**
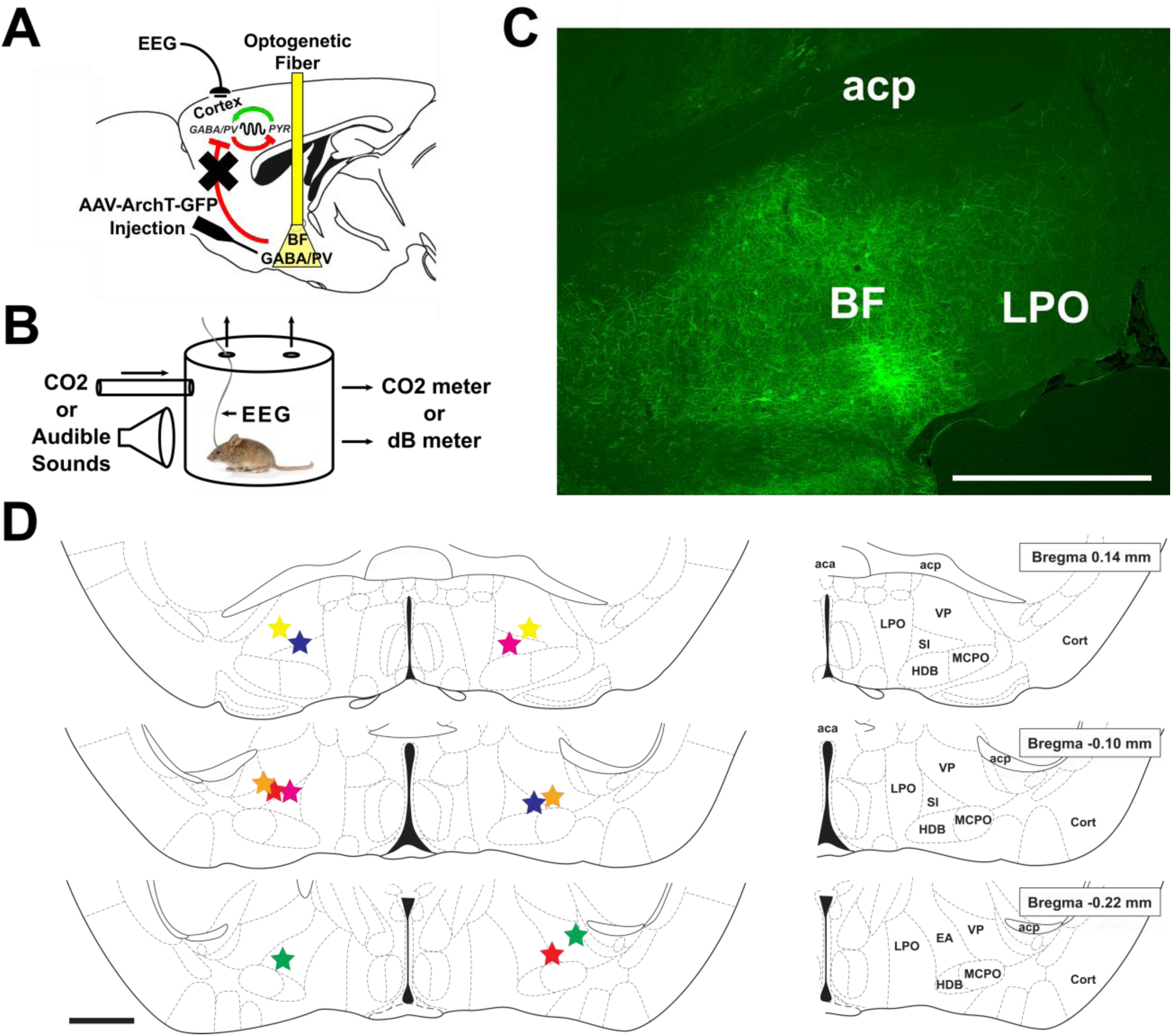
Design of experiments to evaluate the necessity of BF PV neurons in arousals from sleep produced by hypercarbia (excessive CO2) or auditory stimuli. **(A)** Adeno-associated viral vectors with Cre-dependent expression of ArchT-green fluorescent proteins (AAV-Flex-ArchT-GFP) were bilaterally injected into the BF of PV-cre mice, and optogenetic stimulation applied to inhibit arousals produced by exposure to CO2 or sound stimuli. **(B)** Graphic depiction of the housing of the animals and recording of electrophysiology plus CO2 or sound levels. **(C)** Representative photomicrograph example of extensive viral transduction in BF. Scale bar = 1 mm. **(D)** Localization of bilateral optogenetic fiber stimulation sites in BF (N=6). Color coding indicates each individual case. Scale bar = 1mm. For abbreviations, see Figure 2 legend. Atlas templates schematics adapted from Paxinos and Franklin, 2013.

One or more months following surgery, mice were placed in a chamber (**Figure 3B**) where they were exposed to brief (30s) elevations in the ambient level of CO2, cycling every 5 minutes for four hours during the light period. Under baseline conditions, latency to cortical arousal from NREM was reliably induced by increasing CO2 levels from 1 % to 10% inside the chamber (15.99±1.23 s; **Figure 4A**), similar to that reported in Kaur et al., (2013, 2017).

**Figure 4.**
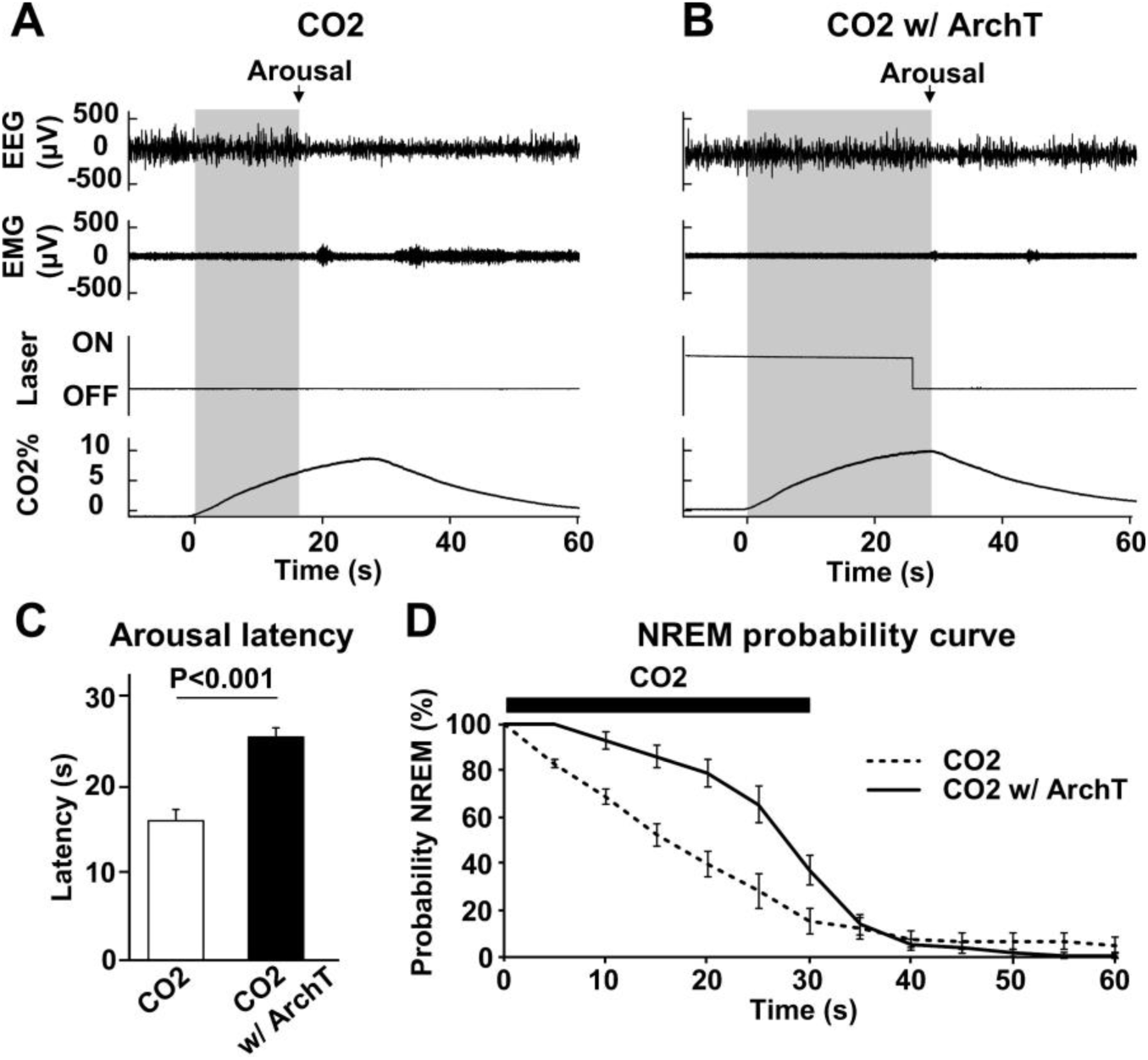
The latency to arouse in response to increased CO2 levels is increased when basal forebrain (BF) parvalbumin (PV) neurons are optogenetically inhibited. **(A)** Representative primary traces of EEG, EMG, laser on/off, and ambient CO2 percentage levels depicted. The grey shaded area shows arousal latency from the start of the stimulus to the point at which the animal wakes, determined as the onset of desynchronized EEG lasting > 2 seconds. **(B)** Representative example in the same animal of increased arousal latency due to BF PV optogenetic inhibition (CO2 w/ ArchT). The grey shaded area is wider, representing a longer arousal latency. **(C)** Mean latencies to arouse from CO2 (N=7) significantly differed, comparing CO2 vs. CO2 w/ ArchT (BF PV inhibition). **(D)** NREM probability curve plots indicate that the probability that animals stay in NREM sleep during CO2 infusion is significantly increased due to BF PV optogenetic inhibition (CO2 w/ ArchT), compared to mock stimulation (CO2).

Transduced BF PV neurons were optically inhibited bilaterally with 40 s of continuous green light (2mW), with laser illuminations10 s preceding CO2 exposure, and continuing during stimuli exposure (30s). When this hypercarbia stimulus was presented while bilaterally inhibiting BF PV neurons, the latency to arousal was markedly increased (ArchT stim; 25.6±1.04 s; **Figure 4B**). Mean data confirmed that latencies were significantly longer when PV+ BF neurons were inhibited (15.99±1.23 s vs. 25.6±1.04 s; t=6.63 (df6), P<0.001)(**Figure 4C**). Probability curve analysis revealed that a greater proportion of late arousals to increasing CO2 occurred with optical inhibition of PV+ BF neurons (**Figure 4D**). The probability of remaining in NREM sleep in the presence of CO2 within the stimulation period was significantly elevated when BF PV neurons were inhibited (f=12.19 (df1), P>0.001).

### Optogenetic inhibition of BF PV neurons inhibits arousals due to auditory stimuli

Unlike parabrachial neurons, which appear to play a preferential role in mediating arousals to hypercarbia (Kaur et al., 2013, 2017), the BF receives multimodal input from many arousal systems (Brown et al., 2012; Zaborszky et al., 2012). Thus, we predicted that BF PV neurons would also be involved in the arousal response to other sensory stimuli. Accordingly, in our next series of experiments, we measured latencies to arousal in response to an auditory stimulus which progressively increased in volume from 0 to 20dB above background over a 30 s period (**Figure 5**). Bilateral optical inhibition of BF PV neurons using ArchT was applied similar to that in hypercarbia experiments (ArchT laser light begins 10 s preceding sound stimuli exposure and continues for 30 s during stimuli exposure; every 5 minutes for four hours). Like hypercarbia stimuli, these auditory stimuli reliably induced arousals from sleep (**Figure 5A**). When sound stimuli were presented while bilaterally inhibiting BF PV neurons, the latency to arousal was significantly increased (**Figures 5A-C**; Mock stim 22.41±0.92 vs. ArchT stim 27.39±1.57 s; paired t-test, t=2.6 (df6), P=0.02). Probability curve analysis revealed a greater proportion of late arousals to auditory stimuli with inhibition of PV+ BF neurons (**Figure 5D**). The probability of remaining in NREM was significantly higher when optical inhibition of BF PV neurons was applied (f=2.81 (df1), P=0.002).

**Figure 5.**
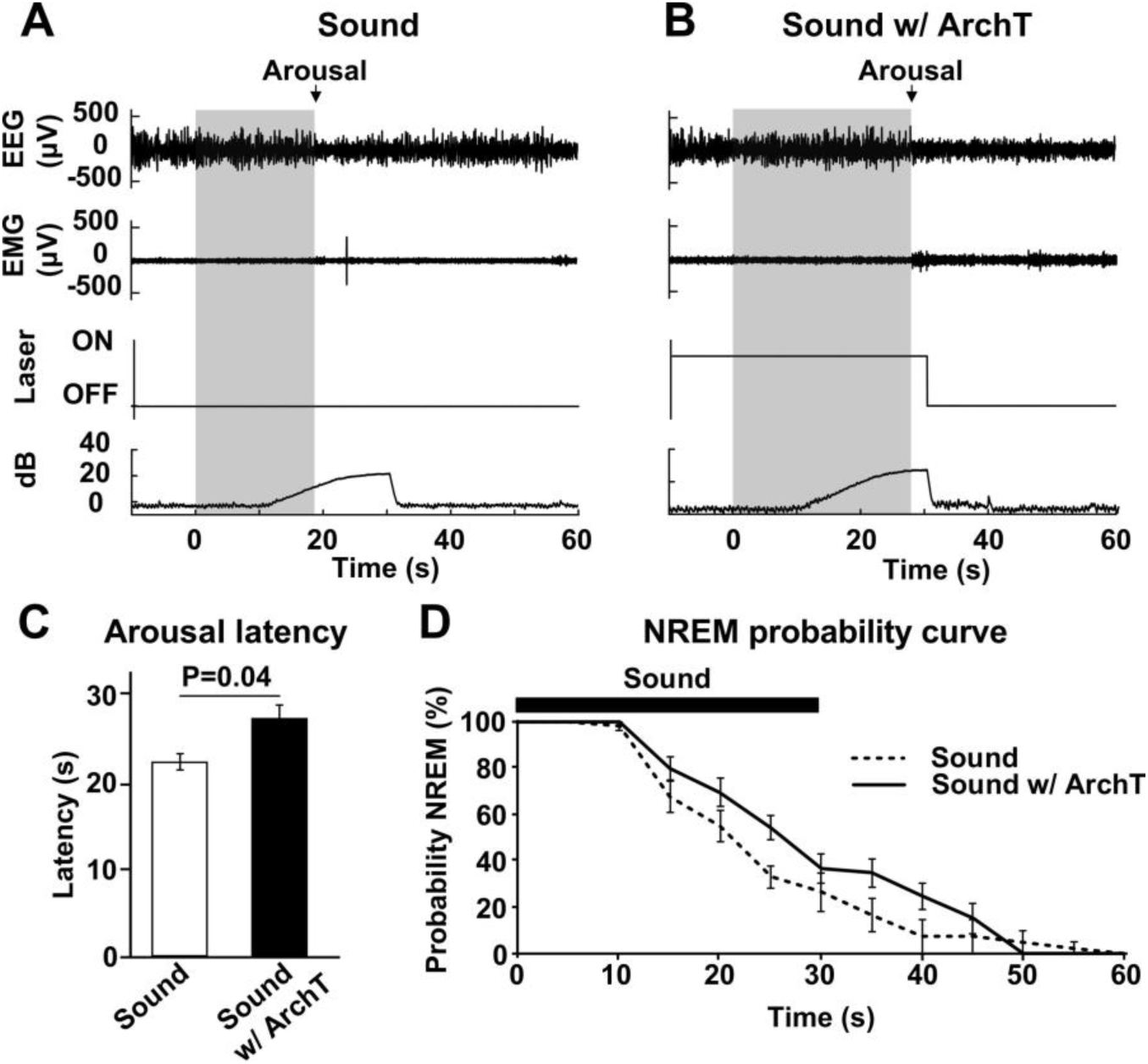
The latency to arouse in response to auditory stimuli is increased when basal forebrain (BF) parvalbumin (PV) neurons are optogenetically inhibited. (A) Representative primary traces of EEG, EMG, laser on/off, and sound decibel levels depicted. The grey shaded area shows arousal latency from the start of the stimulus to the point at which the animal wakes. Representative example in the same animal of increased arousal latency due to BF PV optogenetic inhibition (Sound w/ ArchT). The grey shaded area is wider, representing a longer arousal latency. **(C)** Mean latencies to arouse from sound stimuli exposure (N=7) significantly differed, comparing Sound vs. Sound w/ ArchT (BF PV inhibition). **(D)** NREM probability curve plots indicate that the probability that animals stay in NREM sleep during sound stimuli exposure is significantly increased due to BF PV optogenetic inhibition (Sound w/ ArchT), compared to mock stimulation (Sound). Note: The first 10 secs of sound stimuli were not detectable by the microphone above the background noise.

## DISCUSSION

Physiological systems which mediate arousal from sleep in response to potentially dangerous sensory stimuli are beneficial and important for survival. However, if such arousals occur too frequently or last for longer durations, they can disrupt the restorative aspects of sleep. Such a system must be fast-acting to allow the animal to quickly respond but need not be long lasting. In this study we describe such a role for the cortically projecting BF-PV neurons.

We show that optical stimulations of BF-PV neurons elicit short latency arousals. In contrast optogenetic inhibitions of these neurons increase the latency to arousal in response to internal (hypercarbia) and external (acoustic) sensory signals.

Our optogenetic stimulation experiments here showed that NREM-to-wake arousal latencies were substantially faster than those previously reported for stimulation of cholinergic neurons (∼13.5s, Zant et al., 2016; 25-45 s Irmak and De Lecea 2014) or hypocretin/orexin neurons (20-40 s, Carter et al., 2012). A previous study (Xu et al., 2015) showed that optogenetic stimulation of BF PV neurons could promote transitions to wakefulness but precise measurements of arousal latencies were not reported. Although we show here that BF PV neurons promote rapid arousals, they are not the only ‘rapid arousal system’. For example, optogenetic stimulation of noradrenergic locus coeruleus neurons (< 4s; Carter et al., 2010) or neurotensin neurons in the hypothalamus (< 5s; Naganuma et al., 2019) also produces rapid arousals from sleep.

The short latency to arousal and brief arousal periods observed with BF-PV stimulations parallel clinical sleep apnea data which suggested that cortical arousals are normally brief and sleep fragmentation occurs in the context of relatively minor changes in the total amount of wakefulness (Guilleminault et al., 1976; Dempsey et al., 2010). We observed that repetitive optogenetic stimulation of BF PV neurons for 6 hrs significantly increased the total amounts of wakefulness and decreased amounts of NREM sleep, but these changes were relatively minor (13.6 % increase in wakefulness and 7.2 % decrease in NREM sleep). Furthermore, wake bout durations were significantly increased. These findings are broadly consistent with a previous study which showed that 10 Hz optogenetic stimulation of BF PV neurons promoted wakefulness at the expense of NREM sleep but was less effective compared to optogenetic stimulation of either glutamatergic or cholinergic BF neurons (Xu et al., 2015). In fact, optical stimulation of BF glutamatergic neurons was found to lead to almost constant wakefulness (Xu et al., 2015). Similarly, in a previous study we showed that bilateral BF cholinergic stimulation produced much greater overall amounts of wakefulness (Zant et al., 2016) compared to our present findings. Another heavily studied subpopulation of BF GABAergic neurons, somatostatin-expressing neurons, have been suggested to be sleep-active and promote sleep and are thus unlikely to play a role in rapid-arousal (Xu et al., 2015; Anaclet et al., 2018). Thus, when considering BF neurons, the PV neurons appear to play a more focused role in brief arousals from sleep, more so than other BF populations.

Although gain-of-function experiments are informative, loss-of-function tests are more powerful in testing the role of a particular neurotransmitter system. Our findings indicate that BF PV neurons are integral for arousal from NREM sleep produced by hypercarbia or acoustic stimuli. Recent evidence suggested that the BF is a likely component of the arousal system mediating arousals in response to hypercarbia (Kaur et al., 2017; Kaur and Saper 2019). The BF, along with the lateral hypothalamus and the central nucleus of the amygdala, receives excitatory innervation from the wake promoting PBel (extended lateral aspect of the parabrachial nucleus) area of the brainstem, which is critically involved in arousals to CO2, but not arousals to sounds. Optogenetic inhibition of the majority subpopulation of glutamatergic neurons in PBel, which is characterized by calcitonin gene-related peptide (CGRP) immunoreactivity, delayed latencies to arousal in response to CO2. Interestingly, optogenetic inhibition of transduced PBel CGRP neuronal terminals in BF, LH and CeA delayed arousal latencies, with the most profound effect observed in the BF. This work highlighted the importance of the BF in arousal circuits including the hypercarbia arousal circuit. However, it did not reveal which BF neuronal subgroups are involved.

Ours is the first study to look at any of the subpopulations of wake promoting BF neurons in the response to hypercarbia. We demonstrate for the first time that inhibiting BF PV neurons attenuates arousals from NREM in response to both CO2 and audible sounds. Our results suggest that BF PV neurons are an important component of the ascending arousal circuitry which mediates brief arousals from sleep in response to internal (hypercarbia) or external (auditory) stimuli. Increased activity of BF PV neurons causes very rapid cortical arousals, but they are brief, resulting in relatively minor changes in overall amounts of wakefulness. Conversely, inhibiting BF PV neurons markedly reduces the probability of an arousal from sleep in response to internal or external sensory stimuli. A logical extension of this study suggests BF PV neurons as a candidate target for therapeutic intervention of sleep apnea patients suffering with low arousal thresholds. These neurons may also be an interesting target for treatments to prevent arousals from sleep due to ambient noise.

## AUTHORS’ CONTRIBUTIONS

J.T.M., S.T., J.M.M., J.C.Z., R.E.B., R.W.M., R.E.S., and R.B. conceived and designed the experiments. S.T., J.T.M., D.S.U., C.S., J.C. J.M.M., F.K., and J.C.Z. performed the experiments. S.T., J.T.M., C.S., D.S.U., J.M.M., F.K., M.C.G., and R.B. analyzed the data; J.T.M., R.E.B., J.M.M., R.E.S., and R.B. drafted and revised the manuscript for content. Other authors provided comments on the manuscript. K.D. provided AAV-DIO-ChR2-EYFP through the UNC Vector Core.

## ACKNOWLEDGEMENTS

This work was supported by VA Biomedical Laboratory Research and Development Service Merit Awards I01 BX001404 (RB), I01 BX001356 (REB), I01 BX000270 & I01 BX002774 (RES), I01 BX004500 (JMM) and VA CDA IK2 BX002130 (JMM); and NIH support from R21 NS079866 (RB), R21 NS079866 (REB), R21 NS093000 (REB), R01 MH039683 (REB), T32 HL07901 (JC,FK, DSU) and P01 HL095491 (RES). JTM received partial salary compensation and funding from Merck MISP (Merck Investigator Sponsored Programs) but has no conflict of interest with this work. JTM, JMM, RES, RWM and RB are Research Health Scientists at VA Boston Healthcare System, West Roxbury, MA. The contents of this work do not represent the views of the U.S. Department of Veterans Affairs or the United States Government.

## STAR METHODS

### Animals

Adult (4-6 months) homozygous PV-Cre mice (B6;129P2-*Pvalb^tm1(cre)Arbr^*/J) were purchased from Jackson Laboratory (Stock#008069; Bar Harbor, Maine) and bred in house. Mice were housed at 21°C with a 12-h light/ dark cycle (7:00 AM–7:00 PM), and food and water *ad libitum*. All procedures were performed in accordance with the National Institutes of Health guidelines and in compliance with the animal protocol approved by the VA Boston Healthcare System Institutional Animal Care and Use Committee.

### Viral vector injections

For optical excitation of BF PV neurons we used adeno-associated viral vectors serotype 5 (AAV5), with Cre-recombinase dependent expression of a fusion protein, consisting of ChR2 and EYFP (AAV-DIO-ChR2-EYFP; 3×10^12^ viral particles/ml estimated by DotBlot, University of North Carolina Vector Core; Chapel Hill, NC). For inhibition of BF PV neurons AAV5-CAG-flex-reverse-ArchT-GFP (hereafter referred to as AAV-FLEX-ArchT), bearing the ArchT strain Halorubrum sp. TP009, was purchased from the University of North Carolina vector core. Virion concentration was estimated to be 2 x 10^12^ particles/ml by DotBlot. Viral vectors (500 nL for AAV-DIO-ChR2-EYFP, 1 uL for AAV-FLEX-ArchT), were bilaterally injected into BF (AP 0.0 mm, ML ± 1.6 mm, and ventral -5.0 to -5.3 mm) of PV-Cre mice under isoflurane anesthesia (induction, 5%; maintenance, 1–2%) using a Nanoliter microdispenser (Nanoliter 2010 injector, 23nl/step, slow release; World Precision Instruments; Sarasota, Fl) or a Hamilton syringe (500nl/minute) driven by a high-precision injector pump (model 250; KD Scientific). Following the injections, the scalp incision was sutured closed and mice were allowed to recover. We previously confirmed effective optogenetic excitation and inhibition of BF PV neurons using these vectors *in vitro* (Kim et al., 2015).

### EEG/EMG electrode and fiber optic cannula implantation

Two or more weeks following AAV viral injections, EEG screw electrodes were bilaterally implanted into the skull above the frontal cortices (AP = 1.5-1.9 mm; ML = ±1.0-1.5 mm) under isoflurane anesthesia. A reference screw was placed above the cerebellum, and electromyography (EMG) electrodes were placed in the nuchal muscle. Electrodes were connected to EEG/EMG headmounts (Pinnacle Technology Inc., part # 8402-SS, KS, USA). Fiber optical cannulae (200µm, 0.22 NA optical fiber, Doric Lenses; Quebec City, Quebec, CA) were implanted targeting BF (AP 0.0 mm, ML ±1.6 mm, and ventral 5.0 mm). The headmount and fiber optic cannulae were secured to the skull using dental cement.

### In Vivo EEG/EMG Recordings

For optical excitation work, at least one week following EEG/EMG surgery, mice were tethered and acclimatized to the recording chamber for at least 4 days. After acclimatization, continuous EEG/EMG (8200-K1-SL amplifier; Pinnacle Technology or Model 3600 16-Channel Extracellular Amplifier; A-M Systems) and video recordings were performed before, during, and after optical (or mock control) stimulation of BF PV neurons. The EEG/EMG response to BF light stimulation was recorded using: (1) Sirenia Sleep Pro (Pinnacle Technologies); or (2) the Spike2 analysis program (Cambridge Electronic Design; Cambridge, UK) and a CED1401 analog-to-digital converter (Cambridge Electronic Design). EEG signals were sampled at 2-4 KHz and bandpass filtered at 1–200 Hz. For hypercapnia and acoustic arousal work, mice were tethered and acclimatized to the recording chamber for 4 hours (9am-1pm). After acclimatization, continuous EEG/EMG recordings (8200-K1-SL amplifier; Pinnacle Technology) with and without optical stimulation of BF PV neurons for 4 hours (1pm–5pm). EEG/EMG and other signals were recorded with WinWCP (Strathclyde Institute of Pharmacy and Biomedical Sciences).

### Optical Excitation of BF PV Neurons

Optical stimulation was performed bilaterally to excite BF PV neurons transduced with ChR2 using a fiber-coupled 473-nm solid-state laser diode (Cat # CL473-050-O; 20 mW; CrystaLaser; Reno, Nevada) and an optical fiber (Doric Lenses) connected to the implanted fiber optic cannula with a zirconium sleeve. Software-generated transistor–transistor logic pulses were used to drive the laser light stimulation (Spike2; Cambridge Electronic Design or WinWCP; Strathclyde Institute of Pharmacy and Biomedical Sciences). 10 ms pulses were bilaterally delivered at 40 Hz for 5 s, followed by 55 s of no stimulation. This protocol was repeated for the duration of the experiments (6 hours, ZT 2-8; Figure 1B). Time-of-day matched mock-stimulation (TTL pulses with laser off) in the same mice on a different day served as control.

### Sleep scoring and multi-taper analysis for optogenetic excitation experiments

Vigilance states were scored as one of the following: (1) Wake; active behavior accompanied by desynchronized EEG (low in amplitude), and tonic/phasic motor activity evident in the EMG signal; (2) NREM sleep; more synchronized EEG, higher in amplitude, with particularly notable power in the delta (0.5-4 Hz) band, and low motor activity (EMG); and (3) REM sleep; small amplitude EEG, particularly notable power in the theta (4-9 Hz) band, and phasic motor activity (EMG). Compared to wake, EEG power during REM was significantly reduced in lower delta frequencies (0.5–4 Hz) and increased in the range of theta activity (4–9 Hz).

### Measurement of arousals for optogenetic excitation experiments

For optogenetic stimulations that occurred after at least 10 s of NREM (determined post-hoc during EEG/EMG scoring), the latency to arousal was calculated as the time from the beginning of the stimulation to the end of the last slow wave activity leading to EEG desynchronized activity within 55 s. In < 2% of cases, optogenetic stimulation failed to result in arousal within the next 55s. For such cases, the latency was more than 60 s, extending into the next stimulation period, and not included in evaluations. The median statistic was determined per animal because of the skewed, non-normal distribution of responses. Optogenetic stimulation values were compared with that of the mock-stimulation baseline (BL) day (laser off). We plotted the cumulative probability distribution of remaining in NREM sleep vs. time since stimulation per cycle, averaged for all arousals per case of bilateral optogenetic stimulation (N=14), and compared to all arousals per case of mock (control) stimulation.

### Hypercapnia and auditory stimulation experiments

Mice were recorded in custom built chambers into which CO2 or sounds were delivered (Figure 3C). Ambient CO2 levels were controlled using a three-channel programmable gas mixer (GSM-3; CWE Inc.) at a flow-rate of 1 L/min. CO2 levels where continuously monitored with a cage mounted CO2 sensor (Cozir Sprint). Auditory stimuli were generated using a custom MATLAB script and delivered via WinEDR software (University of Strathclyde, Glasgow, UK) to cage mounted speakers.

Background and stimulus dB were continuously monitored using a cage mounted microphone (MAX4466; Adafruit). Both CO2 and auditory sensor signals were fed via an Arduino Uno to a recording device (USB-6341; National Instruments). From the onset of recordings, EEG/EMG and gas or sound signals were sampled at 1kHz using WinEDR. This software was also used to control switching on/off the lasers, gas and acoustic stimuli. Activation of inhibitory ArchT was conferred by light delivered from a 532 nm laser (MGL-III-532–100mW; Opto Engine, Midvale, UT), delivered to BF via a fiberoptic patch cable (MFP_200/220/900–0.22_2m_FCM-MF1.25; Doric Lenses) into WinEDR.

### Measurement of arousals for hypercapnia and auditory stimulation experiments

For optogenetic stimulations that occurred after at least 10 s of NREM (determined post-hoc during EEG/EMG scoring), the latency to arousal was calculated visually from EEG records as the time from the beginning of the stimuli presentation (CO2 infusion or sound) to the end of the last slow wave activity, indicated by EEG desynchronized activity for >2s. Muscle tone was not a necessary criterion in determining arousals. Values were compared between Mock stim+CO2 vs. optogenetic ArchT BF PV stim+CO2 exposure; or Mock stim+sound stimuli vs. optogenetic ArchT BF PV stim+sound stimuli exposure. We plotted the probability distribution of remaining in NREM sleep vs. time since the beginning of stimulation per cycle, averaged for all arousals per case of ArchT inhibition during CO2 infusion or sound stimuli exposure (laser on), and compared to all arousals per case of non-ArchT (Mock stim laser off+CO2 infusion or sound stimuli) exposure (N=7).

### Perfusion/brain extraction

Mice were anesthetized with sodium pentobarbital (50 mg/ml), exsanguinated with ice-cold PBS, and perfused transcardially with 10% formalin (Cat # HT5011; Sigma-Aldrich; St. Louis, MO). Brains were post-fixed for 2 days in 10% formalin and in 30% sucrose for a third day. 40 µm-thick coronal slices were collected in 1 of 4 wells (phosphate buffered saline) and stored at 4°C.

### Anti-GFP/PV antibody labelling

According to information provided by the manufacturer (RnD Systems; Minneapolis, MN), the sheep anti-PV primary antibody (Cat # AF5058) precipitated a single 12 kDa band on Western blots from rat, mouse, and human brain tissue. This antibody has been previously characterized (e.g., Iijima et al., 2014; Usoskin et al., 2015; Stephany et al., 2016). PV labeling was similar to that previously reported in many brain regions in the mouse (Tamamaki et al., 2003; Xu et al., 2006; Kim et al., 2012; McKenna et al., 2013; Kim et al., 2015) and rat (Endo et al., 1986; Duque et al., 2000; Gritti et al., 2003; Deurveilher et al., 2006), including the cortex, medial septum/vertical limb of the diagonal band, BF, and the thalamic reticular nucleus.

Tissue was first blocked with 0.5% TX-100 in PBS + 3% normal donkey serum for 1 hour. Sections were then incubated overnight in primary antibody (mouse anti-GFP; 1:1000; Cat # MAB3580; EMD Millipore; Billerica, MA) followed by 3 hours of incubation with secondary antibody (donkey anti-mouse IgG conjugated to AlexaFluor 488, green; 1:100; Cat # A-21202; ThermoFisher Scientific; Waltham, MA). This was followed by overnight incubation in the second primary antibody (sheep anti-PV; 1:200; Cat # AF5058; RnD Systems) and then next-day appropriate secondary antibody for 3 hours (donkey anti-sheep IgG conjugated to AlexaFluor 594, red; 1:100; Cat#A-11016; ThermoFisher Scientific). Control stains were performed in slices where primary antibodies were omitted. In some cases, optogenetic fiber optic cannula location was also determined employing cresyl violet staining on another well of tissue, using a previously established protocol (Paxinos and Franklin, 2013).

### Tissue mounting/microscopy and photography

Fluorescently labeled sections were mounted onto gel-alum subbed slides and coverslipped using Vectashield Hard Set mounting medium (Cat # H-1400; Vector Laboratories; Burlingame, CA). Fluorescent microscopy and photography were performed using a Zeiss Image2 microscope, with a Hamamatsu Orca R2 camera (C10600). PV+ cells were identified by red somata (PV stain), co-localized with green (EYFP viral transfected, amplified with anti-GFP antibody) cytoplasm/soma. Labeled cells were quantified using Neurolucida software (Microbrightfield; Williston, VT). Optogenetic fiber optic cannula locations were mapped onto appropriate schematic templates (Franklin and Paxinos, 2008) employing Adobe Illustrator (v.CS5.1). Confocal fluorescence microscopy was performed using a Nikon Eclipse inverted confocal microscope and NIS (Nikon Instruments Software)-element imaging software. One case for BF PV ChR2, and one case for ArchT studies, were not evaluated due to histological complications.

### Data Analysis and Statistics

For optogenetic stimulation experiments (ChR2 and ArchT), comparisons between mock and optogenetic stimulation were performed using paired t-tests. Statistical analysis used SPSS software (release 11.5), and differences were determined to be significant when P<0.05. Cumulative probability distribution analysis (Figures 1E**, 4D, and 5D**) was evaluated using repeated measures ANOVA, based on that previously employed (Kaur et al., 2013, 2017; Zant et al., 2016). Due to inequality of variances (failure of sphericity) found between the two conditions in the optogenetic excitation studies (Figure 1E; Mock vs. ChR2 Stim), data was analyzed using nparLD package in R (R Core Team, 2019) with an LD-F2 (longitudinal data; Condition stratifies Time) design, allowing a robust nonparametric analysis of longitudinal data in factorial settings (Noguchi et al., 2012). Non-parametric probability was represented as the probability that Condition 1 (Mock) is larger than Condition 2 (Stim). All averaged data are shown as means ± standard error.

